# Acquisition of a large virulence plasmid (pINV) promoted temperature-dependent virulence and global dispersal of O96:H19 enteroinvasive *Escherichia coli*

**DOI:** 10.1101/2022.10.27.514154

**Authors:** Sydney L. Miles, Vincenzo Torraca, Zoe A. Dyson, Ana Teresa López-Jiménez, Ebenezer Foster-Nyarko, Claire Jenkins, Kathryn E. Holt, Serge Mostowy

**Affiliations:** Department of Infection Biology, London School of Hygiene and Tropical Medicine, London, United Kingdom; School of Life Sciences, University of Westminster, London, United Kingdom; Gastrointestinal Pathogens and Food Safety (One Health), UK Health Security Agency, United Kingdom; Department of Infectious Diseases, Central Clinical School, Monash University, Melbourne, Victoria, Australia; Wellcome Sanger Institute, Wellcome Genome Campus, Hinxton, United Kingdom

## Abstract

Enteroinvasive *Escherichia coli* (EIEC) and *Shigella* are closely related agents of bacillary dysentery. It is widely viewed that EIEC and *Shigella* species evolved from *E. coli* via independent acquisitions of a large virulence plasmid (pINV) encoding a type three secretion system (T3SS). Sequence Type (ST)99 O96:H19 *E. coli* is an emergent clone of EIEC responsible for recent outbreaks in Europe and South America. Here, we reconstruct the evolutionary history of ST99 *E. coli* using BactDating, revealing distinct phylogenomic clusters of pINV-positive and -negative isolates. To study the impact of pINV acquisition on the virulence of this clone, we developed an EIEC-zebrafish infection model showing that virulence of ST99 EIEC is thermoregulated. Strikingly, zebrafish infection using the oldest available pINV-negative isolate reveals a separate, temperature-independent mechanism of virulence, indicating that ST99 non-EIEC strains were virulent before pINV acquisition. Taken together, these results suggest that an already pathogenic *E. coli* acquired pINV and that virulence of ST99 isolates became thermoregulated once pINV was acquired.

**Importance:** Enteroinvasive *Escherichia coli* (EIEC) and *Shigella* are etiological agents of bacillary dysentery. Sequence Type (ST)99 is an emergent clone of EIEC hypothesised to cause human disease by the recent acquisition of pINV, a large plasmid encoding a type three secretion system (T3SS) that confers the ability to invade human cells. Here, using phylogenomic reconstruction and zebrafish larvae infection, we show that the virulence of ST99 EIEC isolates is highly dependent on temperature, while pINV-negative isolates encode a separate temperature-independent mechanism of virulence. These results highlight that ST99 non-EIEC isolates may have been virulent before pINV acquisition and highlight an important role for pINV acquisition in the emergence of ST99 EIEC in humans, allowing wider dissemination across Europe and South America.

## Introduction

Enteroinvasive *E. coli* (EIEC) and *Shigella* species are Gram-negative, human-adapted pathogens that cause bacillary dysentery. The greatest burden of bacillary dysentery is in low- and middle-income countries (LMICs) (1), although the true burden of EIEC infection is likely underestimated since it is difficult to distinguish from *Shigella*. Historically, *Shigella* was classified as its own genus, with four distinct species, but Multi-Locus Sequence Typing (MLST) and whole-genome sequencing data clearly show *Shigella* spp. are lineages of *E. coli*, as are EIEC (2, 3). Each *Shigella* and EIEC lineage emerged independently within the *E. coli* population, following the horizontal acquisition of a ∼220 kbp virulence plasmid (also known as plasmid of invasion, or pINV) from a currently unknown source (2). pINV encodes a type three secretion system (T3SS) which facilitates the invasion of human epithelial cells and is thermoregulated in both EIEC and *Shigella* (4).

A novel clone of EIEC, of serotype O96:H19 and Multi-Locus Sequence Type (ST) 99, was first described in 2012 in Italy, and has since caused several foodborne outbreaks of moderate to severe diarrheal disease across Europe and South America (5-7). Before 2012, ST99 *E. coli* had not been reported in the literature as causing human disease but had been sporadically isolated from cattle and environmental sources (8). ST99 EIEC isolates have been characterised as possessing the virulence hallmarks of EIEC and *Shigella* (pINV and T3SS) (9), but its metabolic capacity closely resembles that of commensal *E. coli* and it has more recently been associated with *pga*-mediated biofilm formation (6, 9). It has therefore been proposed that ST99 EIEC emerged recently due to the acquisition of pINV.

The zebrafish (*Danio rerio)* larvae model is widely used to study infection biology *in vivo* because of its rapid development and innate immune system which is highly homologous to that of humans (10, 11). Zebrafish have emerged as a valuable vertebrate model to study human enteropathogens like *Shigella* (12), highlighting the key roles of bacterial virulence factors (e.g. T3SS, O-antigen) (13, 14) and cell-autonomous immunity (e.g. autophagy, the septin cytoskeleton) (15, 16) in host-pathogen interactions.

In this Observation, we reconstruct the evolutionary history of ST99 *E. coli* using publicly available whole genome sequences, to understand the role of pINV in its emergence. We develop a temperature-dependent zebrafish infection model to assess the virulence of EIEC and non-EIEC ST99 isolates, highlighting the power of zebrafish infection in studying the evolution of emergent enteropathogens causing disease in humans.

### ST99 EIEC emerged ∼40 years ago

To dissect the evolutionary history of the ST99 clone and its transition to EIEC we analysed all publicly available ST99 genomes (n=92), using the EnteroBase integrated software environment (17). EnteroBase routinely scans short-read archives and retrieves *E. coli* and *Shigella* sequences from the public domain or uses user-uploaded short reads. We used Gubbins (Genealogies Unbiased By recomBinations In Nucleotide Sequences) v.3.2.1 (18) to filter recombinant sites, RaxML v.8.10 to infer a Maximum Likelihood phylogenetic tree, and BactDating v.1.2 (19) to date the phylogeny (Fig 1). Root-to-tip genetic distances were positively associated with year of isolation (R^2^=0.19, p=6×10^−3^), indicating a moderate molecular clock signal to support dating analysis. From this analysis, we estimate the most recent common ancestor (MRCA) of the entire ST99 group existed circa 1776 (95% highest posterior density [HPD], 1360-1927). To test for the presence of pINV, we screened the genomes for *ipa* genes (conserved pINV encoded genes essential for invasion (20)) using Abricate v1.0 (https://github.com/tseemann/abricate). The pINV+ isolates form a distinct cluster, with their MRCA existing circa 1982 (95% HPD, 1965-2011) (Fig 1). This suggests that the ST99 EIEC may have been circulating undetected for ∼30 years before being detected in the 2012 outbreak.

**Figure 1.**
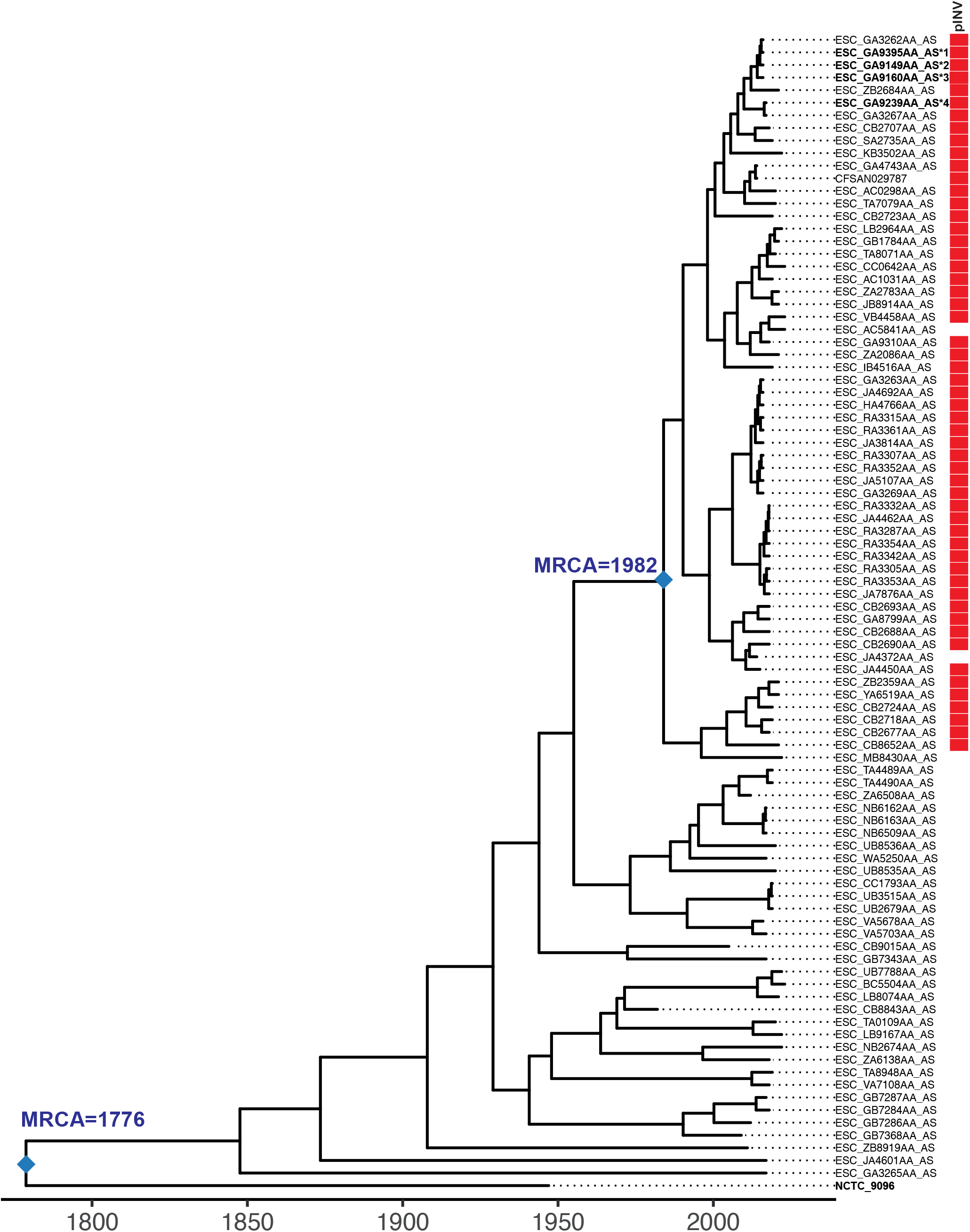
Time calibrated phylogeny of 92 ST99 genomes. BactDating was used to infer a time-calibrated phylogeny, incorporating the output from the recombination detection software, Gubbins. Blue diamonds indicate the internal nodes representing the most recent common ancestors (MRCA) of interest. Tip labels represent assembly barcodes correlating to the isolate accession in Enterobase. Tip labels in bold represent isolates that we tested *in vivo*. The presence of the invasion plasmid, pINV is indicated by a red box in the pINV column. We estimate the MRCA of the whole group to be ∼1776 and ∼1982 for the pINV+ cluster.

To test the role of pINV in the emergence of ST99 EIEC, we selected four recent pINV+ isolates from moderate-to-severe diarrhoeal outbreaks in the UK in 2014 and 2015 (21, 22) to represent contemporary ST99 EIEC, and the oldest available ST99 isolate (∼1945, NCTC 9096, pINV-) to represent ancestral pINV-ST99 (see Fig 1).

### ST99 EIEC virulence is temperature-dependent in zebrafish

The zebrafish infection model has revealed fundamental advances in our understanding of *Shigella* and its ability to infect humans (23). To test the virulence of pINV+ ST99 strains, ∼5000 CFU was injected into the hindbrain ventricle (HBV) of zebrafish larvae at 3 days post fertilisation (dpf) (Fig S1A). Infected zebrafish larvae are typically incubated at 28.5°C for optimal development but we have shown they can also be maintained at 32.5°C (13), allowing the study of temperature-dependent virulence. For the pINV+ strains, we observed ∼75% survival when larvae were incubated at 28.5°C, but only ∼30% survival when incubated at 32.5°C (Fig 2A, Fig S1B-C). In agreement with survival results, CFUs recovered at 6 hours post infection (hpi) were significantly lower at 28.5°C than CFUs recovered at 32.5°C (Fig 2B, Fig S1D-E), suggesting that larvae were more able to control infection at 28.5°C.

**Figure 2.**
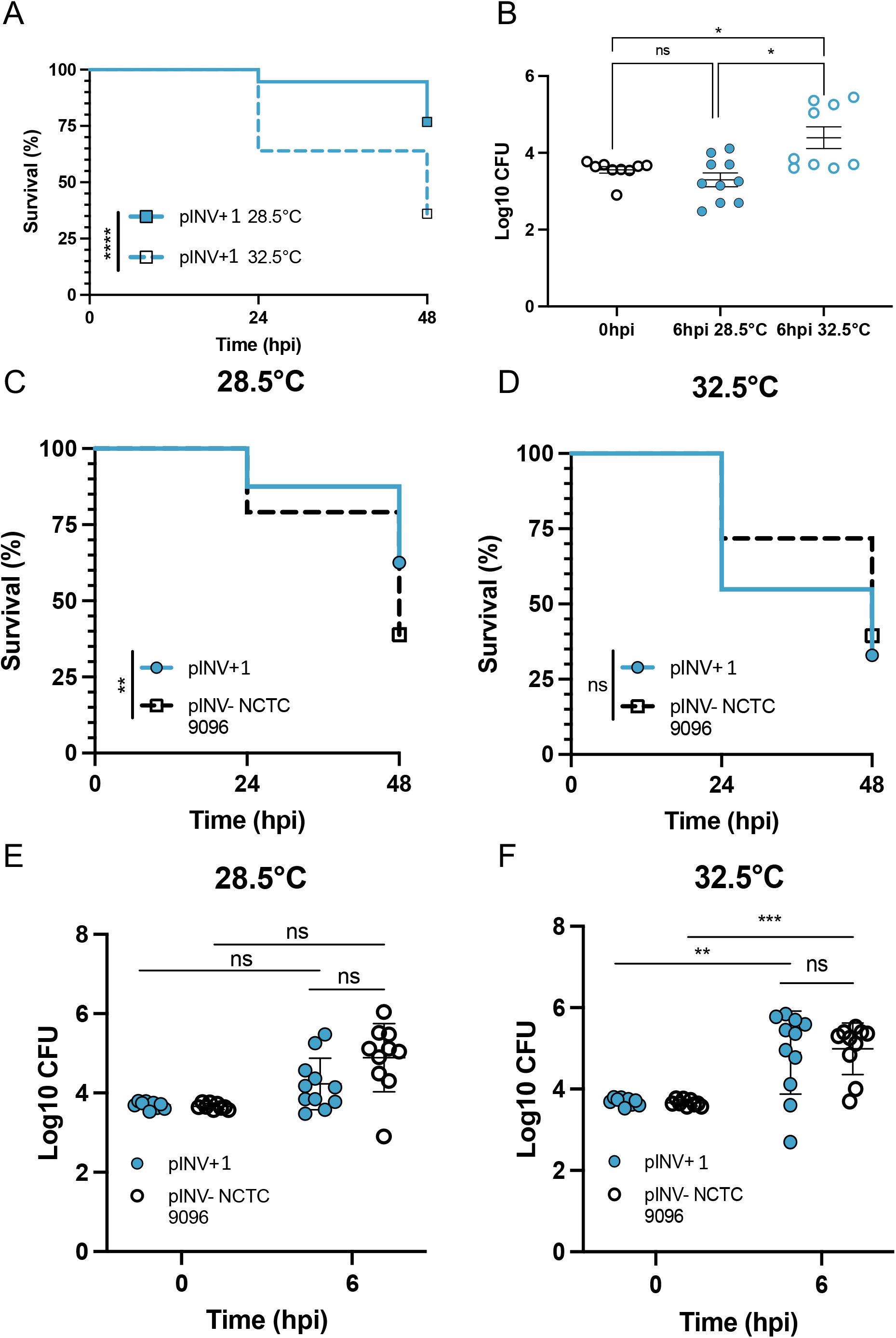
Temperature-dependent and -independent mechanisms of virulence in the ST99 group. Zebrafish larvae at 3 days post fertilisation were injected with 5000 CFU of an ancestral pINV-ST99 strain and a representative pINV+ ST99 strain, before being separated for incubation at 28.5°C or 32.5°C. **A**,**B)** pINV+1 strain exhibits a temperature-dependent virulence with significantly more killing observed at 32.5°C. Enumeration of bacterial burden is also temperature dependent, with greater CFUs quantified at 6 hours post infection from larvae incubated at 32.5°C. Black circles indicate pINV+1 CFUs at 0hpi, blue filled circles indicate pINV+1 CFUs at 6hpi incubated at 28.5°C, blue outlined circles indicate pINV+1 CFUs at 6hpi, at 32.5°C. **C**,**D)** We observe that pINV-strain NCTC 9096 is virulent in the zebrafish model in a non-temperature-dependent manner. Significance tested using Log-rank (Mantel-Cox) test, **p<0.0021. **E**,**F)** Enumeration of CFUs from infected larvae at 6 hpi is temperature dependent in the case of pINV+1, but not pINV-NCTC 9096. *p<0.0332;**p< 0.0021;***p<0.0002;****p<0.0001.

To test if the T3SS in ST99 EIEC is functional and thermoregulated, we compared the secretion of virulence factors by ST99 EIEC and *Shigella flexneri in vitro* (Fig S2). The overall abundance of secreted proteins is lower for ST99 EIEC as compared to *S. flexneri*, but the relative abundance of major secreted effectors appears similar. One exception is SepA, a protein secreted independently of the T3SS, whose presence is known to be variable in other EIEC lineages (24). Although we do not observe significant differences in secretion between 28.5°C and 32.5°C under these *in vitro* conditions tested, the T3SS in ST99 EIEC is clearly thermoregulated (with optimal secretion *in vitro* at 37°C).

Having established a temperature-dependent EIEC-zebrafish infection model, it was next of great interest to test the virulence of an ancestral pINV-ST99 isolate. Since we observed no significant differences in zebrafish survival or bacterial burden between the four pINV+ strains at either 28.5°C or 32.5°C (Fig. S1B-E), we chose one isolate (pINV+1) as a representative pINV+ isolate to compare with the pINV-isolate (NCTC 9096).

### ST99 *E. coli* comprises temperature-dependent and -independent mechanisms of virulence

To test if the virulence of ST99 *E. coli* in zebrafish is dependent on the acquisition of pINV, we compared the virulence of pINV-(NCTC 9096) and pINV+ (pINV+1) strains using our EIEC-zebrafish infection model. Strikingly, NCTC 9096 was significantly more virulent than pINV+1 at 28.5°C, with only ∼35% of infected larvae surviving at 48 hpi (Fig 2C). Although no change in survival of larvae infected with NCTC 9096 is observed at 32.5°C (as compared to that of 28.5°C), survival of pINV+1 infected larvae significantly decrease at 32.5°C, consistent with a role for temperature-dependent virulence. At 32.5°C, we found that both pINV+1 and NCTC 9096 isolates were equally virulent (Fig 2D).

The trend in virulence was also reflected in the quantification of bacterial burden (Fig 2E-F). When incubated at 32.5C, we observed a ∼2 log increase in pINV+1 CFUs enumerated from larvae at 6 hpi, but not when incubated at 28.5°C. We observe a similar increase in NCTC 9096 CFUs quantified, irrespective of temperature. These results clearly show temperature-dependent virulence of the pINV+1 strain, and non-temperature-dependent virulence of the pINV-negative isolate.

## Discussion

It is widely recognised that the acquisition of pINV is a defining feature in the evolution of EIEC and *Shigella* (25). Here, we reconstruct the evolutionary history of ST99 EIEC and propose that a MRCA for the pINV+ group existed in the early 1980s. This suggests that ST99 EIEC may have been circulating undetected for ∼30 years until it was implicated in the 2012 outbreak, perhaps because EIEC infections are typically endemic in regions where surveillance and sequencing of enteropathogens is limited.

We prove that the virulence of ST99 EIEC strains is thermoregulated *in vitro* and *in vivo* (with zebrafish larvae less able to control infection), leading to increased killing and greater bacterial replication at 32.5°C. Some killing is still observed at 28.5°C, suggesting a low-level activation of the T3SS and/or non-T3SS mechanisms of virulence *in vivo*, which would be of interest to test in future studies. These data are consistent with previous reports for pINV-mediated virulence in both *S. flexneri* and *S. sonnei* (4, 13). Our zebrafish infection model highlights the importance of temperature in EIEC virulence and supports the hypothesis that pINV acquisition is the first key step in the evolutionary pathway towards becoming a human-adapted pathogen. Our data using zebrafish infection further shows that non-EIEC ST99 isolates can also cause disease, and that the ability of the ST99 clone to cause disease does not strictly rely on the acquisition of pINV and the transition to EIEC. Considering that pINV-ST99 strain NCTC 9096 is highly virulent *in vivo*, we conclude that it must encode separate, non-thermoregulated mechanism(s) of virulence that becomes less important for human infection once pINV is acquired.

Collectively, our findings illuminate the short evolutionary history of ST99 EIEC and implicate pINV acquisition as a key factor in its emergence and epidemiological success. Our approach also reveals a separate, non-thermoregulated virulence mechanism in a pINV-ST99 isolate, suggesting that an already pathogenic *E. coli* may have acquired pINV. Further studies, including identifying the source of pINV and those isolates likely to acquire it, are important to fully understand and prevent the emergence of novel EIEC and *Shigella* clones infecting humans.

### Methods Bacterial strains

Four pINV+ EIEC strains isolated in diarrhoeal outbreaks from the United Kingdom in 2014/2015 were included in this study (Table 1), strains were identified and sequenced through routine surveillance and kindly shared with us by the UK Health and Security Agency (UKHSA). A pINV-ST99 strain (Table 1) included in this study was obtained from the National Culture Type Collection (NCTC). *Shigella flexneri* M90T was used as a positive control for the *in vitro* secretion assay (26).

**Table 1.**
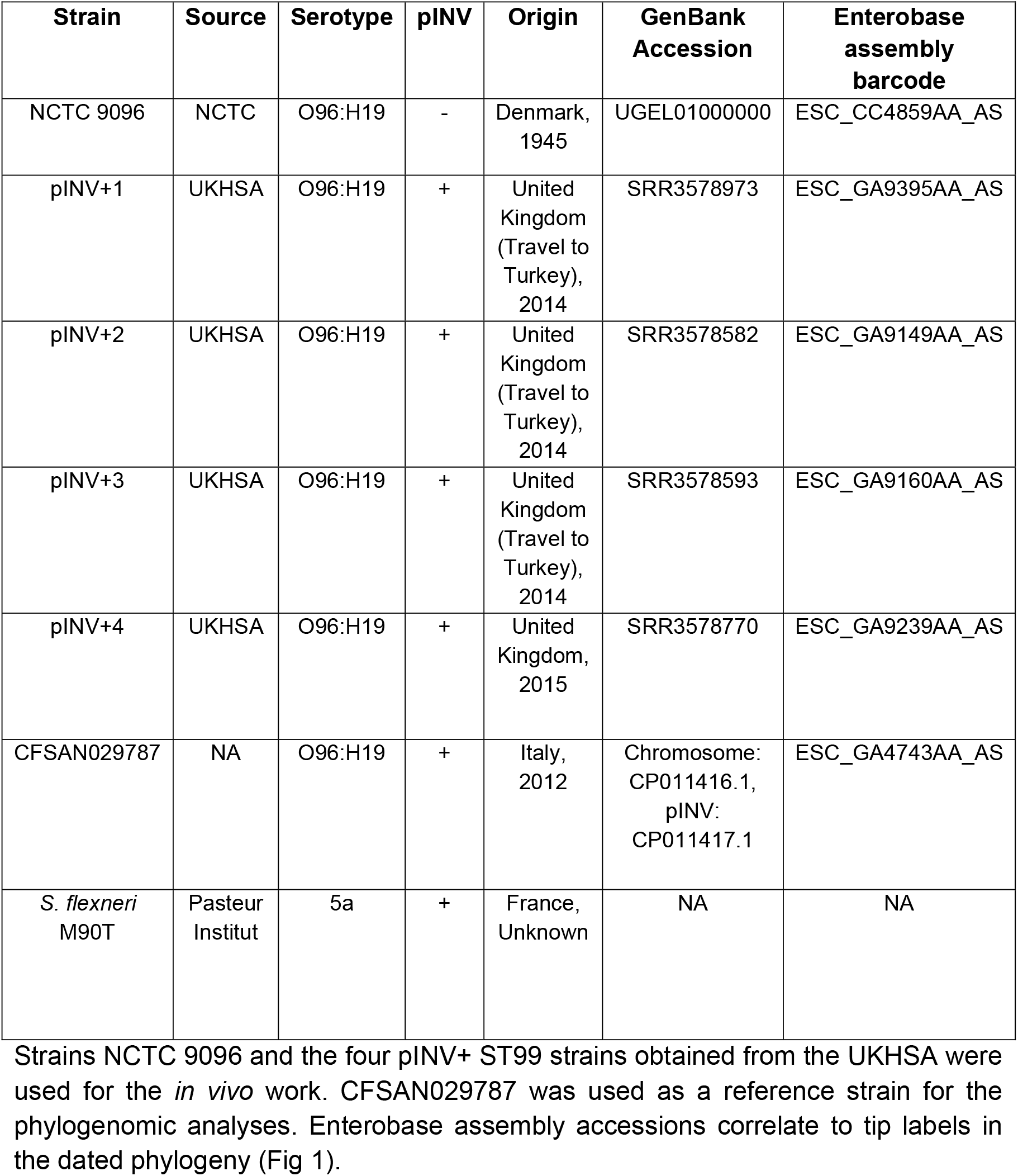
Bacterial strains used in experimental work / as a reference genome.

| Strain | Source | Serotype | pINV | Origin | GenBank Accession | Enterobase assembly barcode |
| --- | --- | --- | --- | --- | --- | --- |
| NCTC 9096 | NCTC | O96:H19 | - | Denmark, 1945 | UGEL01000000 | ESC_CC4859AA_AS |
| pINV+1 | UKHSA | O96:H19 | + | United Kingdom (Travel to Turkey), 2014 | SRR3578973 | ESC_GA9395AA_AS |
| pINV+2 | UKHSA | O96:H19 | + | United Kingdom (Travel to Turkey), 2014 | SRR3578582 | ESC_GA9149AA_AS |
| pINV+3 | UKHSA | O96:H19 | + | United Kingdom (Travel to Turkey), 2014 | SRR3578593 | ESC_GA9160AA_AS |
| pINV+4 | UKHSA | O96:H19 | + | United Kingdom, 2015 | SRR3578770 | ESC_GA9239AA_AS |
| CFSAN029787 | NA | O96:H19 | + | Italy, 2012 | Chromosome: CP011416.1,<br>pINV: CP011417.1 | ESC_GA4743AA_AS |
| *S. flexneri* M90T | Pasteur Institut | 5a | + | France, Unknown | NA | NA |

### Genomic analysis

Enterobase was used to identify publicly available ST99 genomes, using the filter by ST function (17); all ST99 genomes with an associated assembly and isolation date were included in our study (Table S1). Complete genome sequences of strains NCTC 9096 and CFSAN029787 were downloaded from GenBank (accessions UGEL01000000 and CP011416.1, respectively). All genomic analyses were performed using the Cloud Infrastructure for Microbial Bioinformatics (CLIMB) (27). Snippy v.4.6 (https://github.com/tseemann/snippy) was used to generate a core genome alignment, using CFSAN029787 as the reference. Gubbins v.3.2.1 (18) was used to identify and remove recombinant regions of the alignment and RaxML v.8.10 (28) was used to build a maximum likelihood phylogenetic tree, using the General Time Reversible (GTR) GAMMA nucleotide substitution model. BactDating v.1.2 (29) was used to infer the dated phylogeny, using the ‘relaxedgamma’ model and 10^5^ Markov chain Monte Carlo (MCMC) chain iterations. To confirm the temporal signal (association between genetic divergence and time) within our dataset, tip nodes were assigned random dates and the analysis was re-run (this was completed n=10 times). We saw no overlap between the substitution rates of our real data and the randomised datasets (Fig S3). To screen assemblies for the presence of known *E. coli* virulence factors, Abricate (https://github.com/tseemann/abricate) v.1.0 was used, with the e_coli_vf database selected.

### Inoculate preparation

Bacteria were grown on trypticase soy agar (TSA) plates supplemented with 0.01% congo red (Sigma-Aldrich) dye. Single red colonies (pINV+ EIEC) or white colonies (pINV-NCTC 9096) were selected and inoculated into 5mL trypticase soy broth (TSB) and incubated overnight at 37°C, shaking at 400rpm. 400μl of overnight culture was subsequently diluted in 20mL TSB and grown until an optical density of ∼0.6 at 600nm was reached. For zebrafish larvae infections, inoculate preparation was carried out by resuspension of the bacteria at the desired concentration in phosphate buffer saline (PBS, Sigma-Aldrich) containing 2% polyvinylpyrrolidone (Sigma Aldrich) and 0.5% phenol red (Sigma-Aldrich) as previously described (13).

### Ethics statements

Animal experiments were performed according to the Animals (Scientific Procedures) Act 1986 and approved by the Home Office (Project license: P4E664E3C). All experiments were conducted up to 5 days post fertilisation (dpf).

### Zebrafish larvae infection

Wildtype-AB zebrafish embryos were used for *in vivo* studies. Embryos were kept in 0.5x E2 medium supplemented with 0.3μg/mL methylene blue and incubated at 28.5°C unless otherwise stated. Using a microinjector, ∼1nl of bacterial suspension was injected into the hindbrain ventricle (HBV) of 3 days post fertilisation (dpf) zebrafish larvae, following previously described procedures (13). The precise inoculum was determined retrospectively by homogenisation of larvae at 0 hours post infection and plating on TSA plates supplemented with 0.01% Congo red.

For survival assays, zebrafish larvae were visualised using a light stereomicroscope at 24 and 48 hpi, the presence of a heartbeat was used to determine viability. For colony forming unit (CFU) counts, larvae were disrupted in PBS using a pestle pellet blender at 0 and 6 hpi. Serial dilutions in PBS and plating on TSA plates supplemented with 0.01% Congo red were then performed to estimate the bacterial load in each larva. Statistical analysis was performed in GraphPad Prism 9.

#### In vitro secretion assay

Secretion of T3SS effectors was tested as previously described (30). Briefly, bacteria were grown overnight, sub-cultured and grown until exponential phase (OD∼0.4 -0.5) at either 28.5°C, 32.5°C or 37°C. Cultures were then incubated for 3 hours in the presence or absence of Congo red to induce type three secretion. Secreted proteins were collected from culture supernatants, precipitated using trichloroacetic acid (Sigma Aldrich) and then analysed using SDS-PAGE and Coomassie Brilliant Blue R-250 (Bio-Rad) staining.

## Data availability

All genomes used in this study are publicly available in Enterobase, and assembly accessions are provided in Table S1. Gene presence/absence data identified through Abricate for all isolates are provided in Table S2.

### Acknowledgements

We thank members of the Mostowy and Holt labs for helpful discussion. We thank the Biological Services Facility (BSF) at LSHTM for assistance with zebrafish care and breeding. S.L.M. is supported by a Biotechnology and Biological Sciences Research Council LIDo PhD studentship (BB/T008709/1). V.T was supported by an LSHTM/Wellcome Trust Institutional Strategic Support Fund (ISSF) Fellowship (204928/Z/16/Z). A.T.L.J is funded by the European Union’s Horizon 2020 research and innovation program under the Marie Skłodowska – Curie Individual Fellowship (grant agreement No. H2020-MSCA-IF2020-895330). Work in the K.E.H lab is supported by the Bill and Melinda Gates Foundation, Seattle (OPP1175797), KlebNet Project. Work in the S.M. laboratory is supported by a European Research Council Consolidator Grant (grant agreement No. 772853-ENTRAPMENT) and Wellcome Trust Senior Research Fellowship (206444/Z/17/Z).

## Figure legends

**Figure S1. Virulence of ST99 EIEC isolates is temperature dependent**. **A)** Schematic of a zebrafish larva, indicating the hindbrain ventricle as the infection site. **B-E)** 3-day post fertilisation larvae were injected with 5000 CFU of bacteria before being separated for incubation at 28.5°C or 32.5°C. **B**,**C)** pINV+ ST99 strains exhibit a temperature-dependent virulence, with significantly more killing observed at 32.5°C. No differences in survival were observed between strains (p= 0.22). Significance was tested using Log-rank (Mantel-Cox) test. **D**,**E**). Enumeration of bacterial burden is also temperature dependent in pINV+ strains, with greater CFUs quantified at 6 hours post infection from larvae incubated at 32.5°C.

**Figure S2. *S*ecretion of virulence factors *in vitro* by ST99 EIEC pINV+1**. Low exposure (top) and high exposure (bottom) of SDS-PAGE gel stained with Coomassie blue, showing secreted factors by ST99 EIEC pINV+1 in presence or absence of congo red, at different temperatures. *S. flexneri* M90T is used as a positive control. Several well characterised secreted virulence factors are identified and labelled based on their abundance and molecular weight.

**Figure S3. Results of date randomisation test to test for temporal signal within our dataset**. To test for temporal signal, isolation dates were randomised n=10 times and the analysis was re-run (R1-R10). We saw no overlap with the substitution rate of our real data, indicating the suitability of our dataset for performing a dated phylogenomic analysis.

**Table S1. Details and metadata for all genomes downloaded from Enterobase**. The assembly barcode is used to identify genomes in the phylogenetic tree (Fig 1).

**Table S2. Gene presence / absence data for all virulence factors identified using Abricate**. 1 indicates presence and 0 indicates absence. Presence of *ipa* genes were used to assess pINV presence.

## References

1. Kotloff KL, Nataro JP, Blackwelder WC, Nasrin D, Farag TH, Panchalingam S, Wu Y, Sow SO, Sur D, Breiman RF, Faruque ASG, Zaidi AKM, Saha D, Alonso PL, Tamboura B, Sanogo D, Onwuchekwa U, Manna B, Ramamurthy T, Kanungo S, Ochieng JB, Omore R, Oundo JO, Hossain A, Das SK, Ahmed S, Qureshi S, Quadri F, Adegbola RA, Antonio M, Hossain MJ, Akinsola A, Mandomando I, Nhampossa T, Acácio S, Biswas K, O’Reilly CE, Mintz ED, Berkeley LY, Muhsen K, Sommerfelt H, Robins-Browne RM, Levine MM. 2013. Burden and aetiology of diarrhoeal disease in infants and young children in developing countries (the Global Enteric Multicenter Study, GEMS): a prospective, case-control study. The Lancet 382:209–222.

2. Pupo GM, Lan R, Reeves PR. 2000. Multiple independent origins of Shigella clones of Escherichia coli and convergent evolution of many of their characteristics. Proceedings of the National Academy of Sciences 97:10567–10572.

3. Sahl Jason W, Morris Carolyn R, Emberger J, Fraser Claire M, Ochieng John B, Juma J, Fields B, Breiman Robert F, Gilmour M, Nataro James P, Rasko David A. 2015. Defining the Phylogenomics of Shigella Species: a Pathway to Diagnostics. Journal of Clinical Microbiology 53:951–960.

4. Falconi M, Colonna B, Prosseda G, Micheli G, Gualerzi CO. 1998. Thermoregulation of Shigella and Escherichia coli EIEC pathogenicity. A temperature-dependent structural transition of DNA modulates accessibility of virF promoter to transcriptional repressor H-NS. The EMBO journal 17:7033–7043.

5. Escher M, Scavia G, Morabito S, Tozzoli R, Maugliani A, Cantoni S, Fracchia S, Bettati A, Casa R, Gesu GP, Torresani E, Caprioli A. 2014. A severe foodborne outbreak of diarrhoea linked to a canteen in Italy caused by enteroinvasive Escherichia coli, an uncommon agent. Epidemiology and Infection 142:2559–2566.

6. Iqbal J, Malviya N, Gaddy Jennifer A, Zhang C, Seier Andrew J, Haley Kathryn P, Doster Ryan S, Farfán-García Ana E, Gómez-Duarte Oscar G. 2022. Enteroinvasive Escherichia coli O96:H19 is an Emergent Biofilm-Forming Pathogen. Journal of Bacteriology 204:e00562–21.

7. Newitt S, MacGregor V, Robbins V, Bayliss L, Chattaway MA, Dallman T, Ready D, Aird H, Puleston R, Hawker J. 2016. Two Linked Enteroinvasive Escherichia coli Outbreaks, Nottingham, UK, June 2014. Emerging infectious diseases 22:1178–1184.

8. Bai X, Scheutz F, Dahlgren HM, Hedenström I, Jernberg C. 2021. Characterization of Clinical Escherichia coli Strains Producing a Novel Shiga Toxin 2 Subtype in Sweden and Denmark. Microorganisms 9:2374.

9. Michelacci V, Prosseda G, Maugliani A, Tozzoli R, Sanchez S, Herrera-León S, Dallman T, Jenkins C, Caprioli A, Morabito S. 2016. Characterization of an emergent clone of enteroinvasive Escherichia coli circulating in Europe.

10. Howe K, Clark MD, Torroja CF, Torrance J, Berthelot C, Muffato M, Collins JE, Humphray S, McLaren K, Matthews L, McLaren S, Sealy I, Caccamo M, Churcher C, Scott C, Barrett JC, Koch R, Rauch G-J, White S, Chow W, Kilian B, Quintais LT, Guerra-Assunção JA, Zhou Y, Gu Y, Yen J, Vogel J-H, Eyre T, Redmond S, Banerjee R, Chi J, Fu B, Langley E, Maguire SF, Laird GK, Lloyd D, Kenyon E, Donaldson S, Sehra H, Almeida-King J, Loveland J, Trevanion S, Jones M, Quail M, Willey D, Hunt A, Burton J, Sims S, McLay K, Plumb B, et al. 2013. The zebrafish reference genome sequence and its relationship to the human genome. Nature 496:498–503.

11. Gomes MC, Mostowy S. 2020. The Case for Modeling Human Infection in Zebrafish. Trends in Microbiology 28:10–18.

12. Mostowy S, Boucontet L Fau-Mazon Moya MJ, Mazon Moya Mj Fau-Sirianni A, Sirianni A Fau-Boudinot P, Boudinot P Fau-Hollinshead M, Hollinshead M Fau-Cossart P, Cossart P Fau-Herbomel P, Herbomel P Fau-Levraud J-P, Levraud Jp Fau-Colucci-Guyon E, Colucci-Guyon E. 2013. The zebrafish as a new model for the in vivo study of Shigella flexneri interaction with phagocytes and bacterial autophagy.

13. Torraca VA-O, Kaforou MA-O, Watson JA-O, Duggan GM, Guerrero-Gutierrez HA-O, Krokowski S, Hollinshead M, Clarke TB, Mostowy RA-O, Tomlinson GS, Sancho-Shimizu VA-O, Clements AA-OX, Mostowy SA-O. 2019. Shigella sonnei infection of zebrafish reveals that O-antigen mediates neutrophil tolerance and dysentery incidence.

14. Willis A R., Torraca V, Gomes Margarida C, Shelley J, Mazon-Moya M, Filloux A, Lo Celso C, Mostowy S. 2018. Shigella-Induced Emergency Granulopoiesis Protects Zebrafish Larvae from Secondary Infection. mBio 9:e00933–18.

15. Van Ngo H, Robertin S, Brokatzky D, Bielecka MK, Lobato-Márquez D, Torraca V, Mostowy S. 2022. Septins promote caspase activity and coordinate mitochondrial apoptosis. Cytoskeleton.

16. Mostowy S, Boucontet L, Mazon Moya MJ, Sirianni A, Boudinot P, Hollinshead M, Cossart P, Herbomel P, Levraud J-P, Colucci-Guyon E. 2013. The Zebrafish as a New Model for the In Vivo Study of Shigella flexneri Interaction with Phagocytes and Bacterial Autophagy. PLOS Pathogens 9:e1003588.

17. Zhou Z, Alikhan N-F, Mohamed K, Fan Y, Achtman M. 2020. The EnteroBase user’s guide, with case studies on Salmonella transmissions, Yersinia pestis phylogeny, and Escherichia core genomic diversity.

18. Croucher NJ, Page AJ, Connor TR, Delaney AJ, Keane JA, Bentley SD, Parkhill J, Harris SR. 2015. Rapid phylogenetic analysis of large samples of recombinant bacterial whole genome sequences using Gubbins. Nucleic Acids Research 43:e15–e15.

19. Didelot X, Croucher NJ, Bentley SD, Harris SR, Wilson DJ. 2018. Bayesian inference of ancestral dates on bacterial phylogenetic trees.

20. Venkatesan MM, Buysse JM, Kopecko DJ. 1989. Use of Shigella flexneri ipaC and ipaH gene sequences for the general identification of Shigella spp. and enteroinvasive Escherichia coli. J Clin Microbiol 27:2687–91.

21. Michelacci V, Tozzoli R, Arancia S, D’Angelo A, Boni A, Knijn A, Prosseda G, Greig DR, Jenkins C, Camou T, Sirok A, Navarro A, Schelotto F, Varela G, Morabito S. 2020. Tracing Back the Evolutionary Route of Enteroinvasive Escherichia coli (EIEC) and Shigella Through the Example of the Highly Pathogenic O96:H19 EIEC Clone. Frontiers in cellular and infection microbiology 10:260–260.

22. Cowley LA, Oresegun DR, Chattaway MA, Dallman TJ, Jenkins C. 2018. Phylogenetic comparison of enteroinvasive Escherichia coli isolated from cases of diarrhoeal disease in England, 2005–2016. Journal of Medical Microbiology 67:884–888.

23. Duggan GM, Mostowy S. 2018. Use of zebrafish to study Shigella infection.

24. Lan R, Alles MC, Donohoe K, Martinez MB, Reeves PR. 2004. Molecular evolutionary relationships of enteroinvasive Escherichia coli and Shigella spp. Infection and immunity 72:5080–5088.

25. Pasqua M, Michelacci V, Di Martino ML, Tozzoli R, Grossi M, Colonna B, Morabito S, Prosseda G. 2017. The Intriguing Evolutionary Journey of Enteroinvasive E. coli (EIEC) toward Pathogenicity.

26. Sansonetti PJ, Kopecko DJ, Formal SB. 1982. Involvement of a plasmid in the invasive ability of Shigella flexneri. Infect Immun 35:852–60.

27. Connor TR, Loman NJ, Thompson S, Smith A, Southgate J, Poplawski R, Bull MJ, Richardson E, Ismail M, Thompson SE, Kitchen C, Guest M, Bakke M, Sheppard SK, Pallen MJ. 2016. CLIMB (the Cloud Infrastructure for Microbial Bioinformatics): an online resource for the medical microbiology community.

28. Stamatakis A. 2014. RAxML version 8: a tool for phylogenetic analysis and post-analysis of large phylogenies. Bioinformatics 30:1312–3.

29. Didelot X, Croucher NJ, Bentley SD, Harris SR, Wilson DJ. 2018. Bayesian inference of ancestral dates on bacterial phylogenetic trees. Nucleic Acids Research 46:e134–e134.

30. Reinhardt J, Kolbe M. 2014. Secretion Assay in Shigella flexneri. Bio-protocol 4:e1302.

